# Draft genome of a porcupinefish, *Diodon Holocanthus*

**DOI:** 10.1101/775387

**Authors:** Mengyang Xu, Xiaoshan Su, Mengqi Zhang, Ming Li, Xiaoyun Huang, Guangyi Fan, Xin Liu, He Zhang

## Abstract

The long-spine porcupinefish, *Diodon holocanthus* (Diodontidae, Tetraodontiformes, Actinopterygii), also known as the freckled porcupinefish, attracts great interest of ecology and economy. Its distinct characteristics including inflation reaction, spiny skin and tetradotoxin, however, have not been fully studied without a complete genome assembly.

In this study, the whole genome of a single individual was sequenced using single tube-Long Fragment Read co-barcode reads, generating 154.3 Gb of paired-end data (219.8× depth). The gap was further filled using small amount of Oxford Nanopore MinION long read dataset (11.4Gb, 15.9× depth). Taking full use of long, medium, short-range of genome assembly information, the final assembled sequences with a total length of 650.02 Mb obtained contig and scaffold N50 sizes of 2.15 Mb and 8.13 Mb, respectively, despite of high repetitive content. Benchmarking Universal Single-Copy Orthologs captured 95.7% (2,474) of core genes to assess the completeness. In addition, 206.5 Mb (32.10%) of repetitive sequences were identified, and 20,840 protein-coding genes were annotated, among which 18,281 (87.72%) proteins were assigned with possible functions.

This is the first demonstration of *de novo* genome of the porcupinefish, which will benefit downstream analysis of ontogeny, phylogeny, and evolution, and improve the exploration of its unique defensive mechanism.

## I Introduction

Pufferfish are small vertebrate fish in the order Tetraodontiformes. As indicated by their name, most pufferfish can inflate themselves, obviously expanding in size in an effort to fend off predators. This inflation ability is found in all members of the sister taxa Tetraodontidae and Diodontidae ^1,2^. A small number of pufferfish species, porcupinefish, outward spines, as an additional defensive adaptation. Among them, *Diodon holocanthus* is widely distribute in the tropical and warm waters of the Pacific, the Atlantic, and the Indian Ocean. *D. holocanthus* is a demersal fish species which can reproduce several times a year ^3^ and is of significant economic importance.

The inflation mechanism, genome size variation, and genome evolution of pufferfish has been investigated ^1,2,4,5^. The interrelationships and phylogeny of porcupinefish has been studied using single nuclear-encoded genes (e.g. *RAG1*) or mitogenomes ^6–8^. However, the lack of a sequenced whole genome hampered the attempts to explore the key role in adaption and evolution as well as its unique skin character and defensive mechanism.

In this study, we sequenced *Diodon holocanthus* genome using single tube-Long Fragment Read co-barcoded (stLFR) technology, reported the assembly, repeat and gene annotation for the first porcupinefish. The final draft genome assembly was approximately 650.02 Mb with a contig N50 of 2.15 Mb and scaffold N50 of 8.13 Mb. A total of 20,840 protein-coding genes were predicted from the genome assembly. The determination of genomic resource of the porcupinefish will be of significance to improve the understanding of its unique morphological and physiological characteristics.

## II Methods

### Sample collection, library construction and genome sequencing

A single *Diodon holocanthus* fish (Figure 1) was purchased from a seafood market at Xiamen, Fujian province, southeast China. Genomic DNA was extracted from the muscle tissue using a conventional method for high molecular weight DNA ^9^ (liquid nitrogen grinding, followed by phenol-chloroform-Isoamyl alcohol extraction and ethanol precipitation). A stLFR library ^10^ was prepared by using the MGIEasy stLFR Library Prep kit (PN: 1000005622) and sequenced on BGISEQ-500 platform.

**Figure 1.**
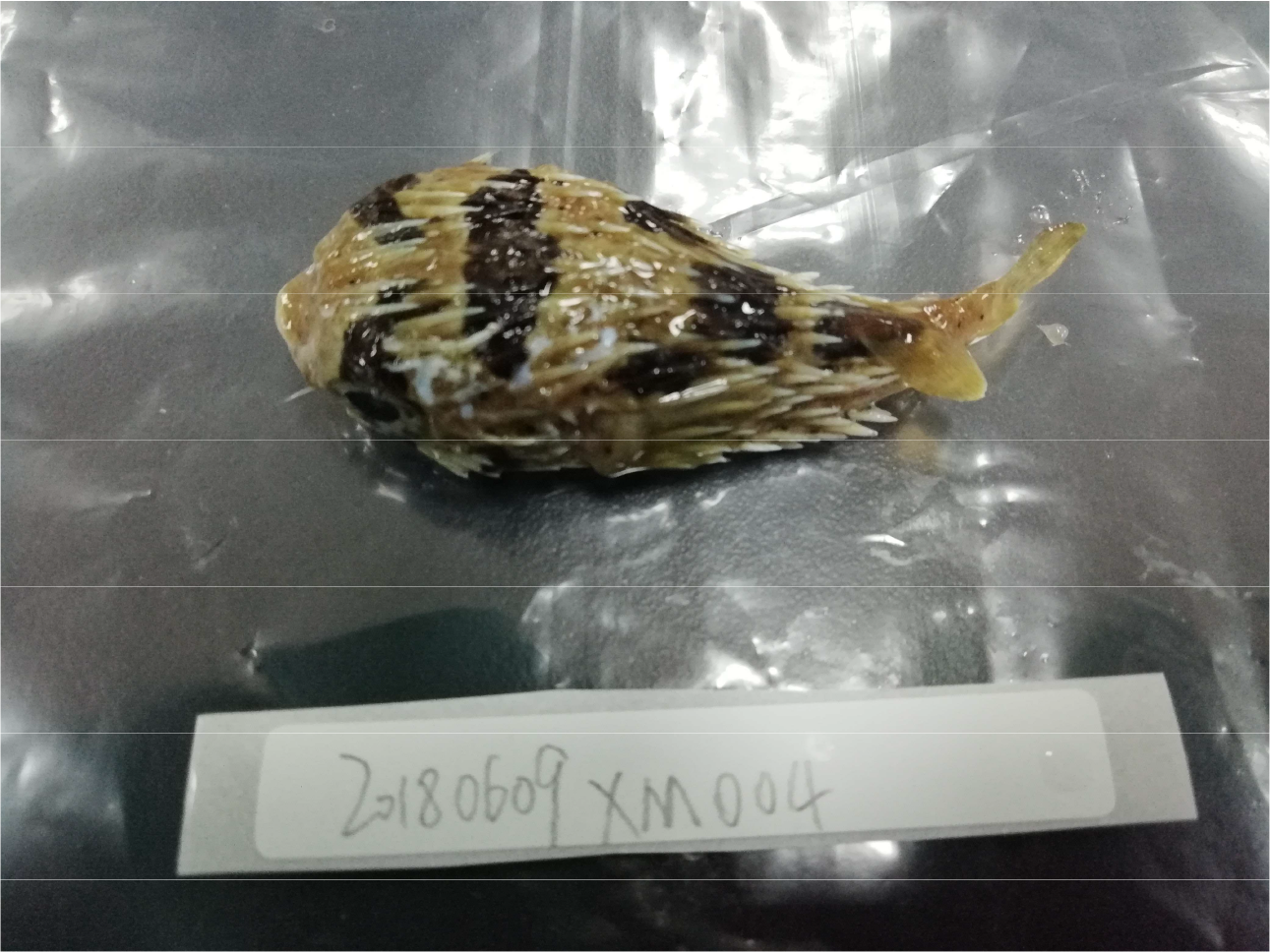
Photograph of *Diodon holocanthus*. (*Credit: M.Z*.)

A total of 637.6 million 100bp paired-end stLFR reads were generated, representing 154.3 Gb of nucleotide sequences, with 77.4% bases ≥ Q30. Raw data was filtered using SOAPfilter software (version 2.2) ^11^ to obtain 93.6 Gb clean reads excluding the adaptor sequences, reads containing more than 10% unidentified nucleotides, and low-quality reads containing more than 50% bases with Phred quality score ≤10, and reads whose sequences are exactly identical, that is, PCR duplications.

A total of 11.4 Gb of sequence data, representing 15.9× theoretical coverage, were produced using the MinION (Oxford Nanopore Technologies,ONT) nanopore sequencer. The average and N50 read length are 17.3Kb and 23.6Kb, respectively.

### Genome features revealed by k-mer analysis

The high-quality clean reads from the short-insert reads (250 bp) were used to estimate the genomic information of *Diodon holocanthus* and 17-mer frequency information was generated based on *k*-mer analysis using KMERFREQ_AR (version 2.0.4) ^11^ for estimation of genome size, heterozygosity, and repeat content. The calculated results provided by the software GEC (version 1.0.0) ^12^ show that the estimated haploid genome size was 701.96 Mb, with a repeat content of 36.35% and a heterozygosity level of 0.76% represented in the first peak of the distribution (Figure 2). The high level of repeat content indicated a troublesome genome characteristic for *de novo* assembly.

**Figure 2.**
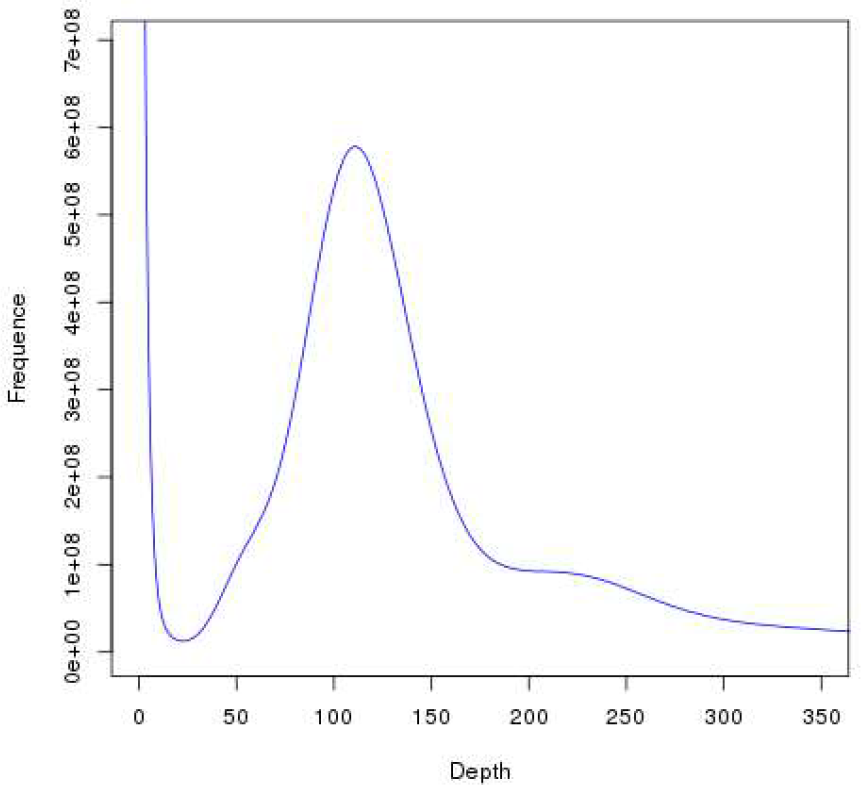
Distribution of 17-mer frequency. In total 93.6 Gb of high-quality short insert reads (250 bp) were used to generate the 17-mer depth distribution curve frequency information.

**Figure 3.**
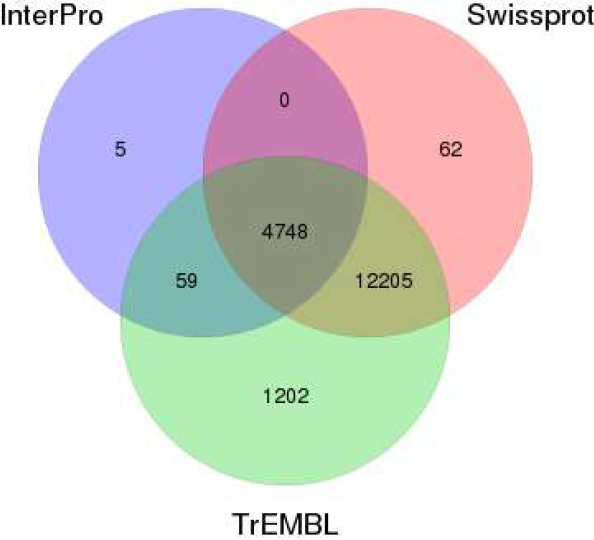
Venn diagram of the number of genes with functional annotation using multiple public databases.

### *De novo* genome assembly

The stLFR data were split into two parts: paired-end 100bp short reads and their corresponding barcode information. Each series of 40-base barcode sequences refers to a unique stLFR barcode ID, a combination of 3 unique tube indices.

Supernova (version 2.1.1) ^13^ is a well-established *de novo* assembler based on a de Bruijn graph algorithm originally designed for barcoded data from 10X Genomics Chromium Linked-Reads. To allow the use of Supernova for stLFR sequencing reads, stLFR barcodes (over 10 million) were parsed (transformed) to generate barcodes compatible with the 10X Genomics format (4.7 million). Scripts are available on GitHub (https://github.com/BGI-Qingdao/stlfr2supernova_pipeline).

We assembled the *Diodon holocanthus* genome, using stLFR clean reads with transformed barcodes, with Supernova as follows: build a 48-mer DBG based on shared *k*-mers, map barcodes to the de Bruijn graph, use barcode information to scaffold, partition the graph and make local assembly, phase based on barcode information, close gap, and reuse barcode and copy number information to further scaffold. The Supernova assembled a genome with scaffold N50 of 6,098,089 bp and contig N50 of 70,841 bp.

To make full use of the diversity of the stLFR barcode information, SLR-superscaffolder (Version 0.9.0) ^14^ was further applied to improve the scaffolds. First, the short reads were aligned back to the Supernova draft scaffolds using BWA-MEM (version 0.7.17) ^15^, in order to distribute the barcode information to scaffolds. Then, according to the shared barcodes, scaffolds were further gathered together and clustered in longer super-scaffolds. In each scaffold group, the order of scaffolds was determined based on a Mean-Spanning-Tree (MST) algorithm, while the orientation of each scaffold was chosen by the Jaccard similarity score with its nth-order neighbors. Contigs that did not belong to any draft scaffolds were also filled in the super-scaffolds based on both barcode and PE information. The scaffold N50 and N90 were improved from 6,098,089 and 976,586 bp to 8,075,516 and 1,206,047 bp, respectively. Note that the re-scaffolding procedure had no effect on contig sequences, only orientation. GapCloser (Version 1.12) ^11^ were applied to reduce gap regions. Considering all contigs, this step enhanced the contig N50 and N90 sizes to 143,286 and 36,130 bp, respectively. Considering the high repetitive content, approximately 15.9× Oxford Nanopore MinION reads were used to overcome the gaps induced by repeats, which was impossible by short reads and kmer extension algorithm, thus further improving assembly quality by TGSGapFiller (Unpublished, available on GitHub https://github.com/BGI-Qingdao/TGSGapFiller). Using such low coverage of long reads, the final assembly of the porcupinefish genome reached a total length of 650.02 Mb, which is similar to the estimated size, with contig N50 of 2,149,931bp, and scaffold N50 of 8,129,349 bp.

Compared with other recently published tetraodontiforme genomes, the total size is almost twice of them, and even larger than two relative species, which is consistent with previous work ^5^. Using stLFR dataset, along with only 15.9× ONT long reads, the contig N50 is comparable to that of those using massive expensive third-generation long reads although the scaffold N50 is relatively smaller due to the lack of BioNano or HiC long-range information ^16^. Both contig and scaffold N50 are drastically larger than that of genomes previously assembled by short reads ^17,18^.

To assess the quality of the genome assembly, we performed the Benchmarking Universal Single-Copy Orthologs (BUSCO) software (Version 3.0.2) ^19^. The genome was queried (options ‘–m geno –sp zebrafish’) against the “metazoa.odb9” lineage set containing 65 eukaryotic organisms to assess the coverage of core eukaryotic genes. Using the vertebrata_odb9 database, BUSCO analysis revealed that 95.7% (91.5% single-copy genes and 4.2% duplicates) of the expected vertebrate genes were complete.

## III Results and Discussions

### Repeat Content

Repetitive sequences usually refer to two major types of repetitive sequences: tandem repeats and interspersed repeats. For the repeat annotation of the porcupinefish genome, both homology-based predictions and *de novo* methods were employed. In the homolog-based methods, RepeatMasker and ProteinMask (version 3.2.9) ^20^ were utilized to detect interspersed repeats searching, against the published RepBase 16.02 ^21^ sequences. In the *de novo* method, the interspersed repeats in the genome were identified using RepeatMasker and RepeatModeler (version 1.1.0.4) ^22^. Tandem Repeats Finder (TRF version 4.04) ^23^ was subsequently used to search for tandem repeats. A total of 211 Mb of non-redundant repetitive sequences in were identified in the porcupinefish genome, accounting for 32,91% of the whole genome (Table 2). The most predominant elements were long interspersed nuclear elements (LINEs), which accounted for 15.17% (97.6 Mb) of the genome (Table 3).

**Table 1.**
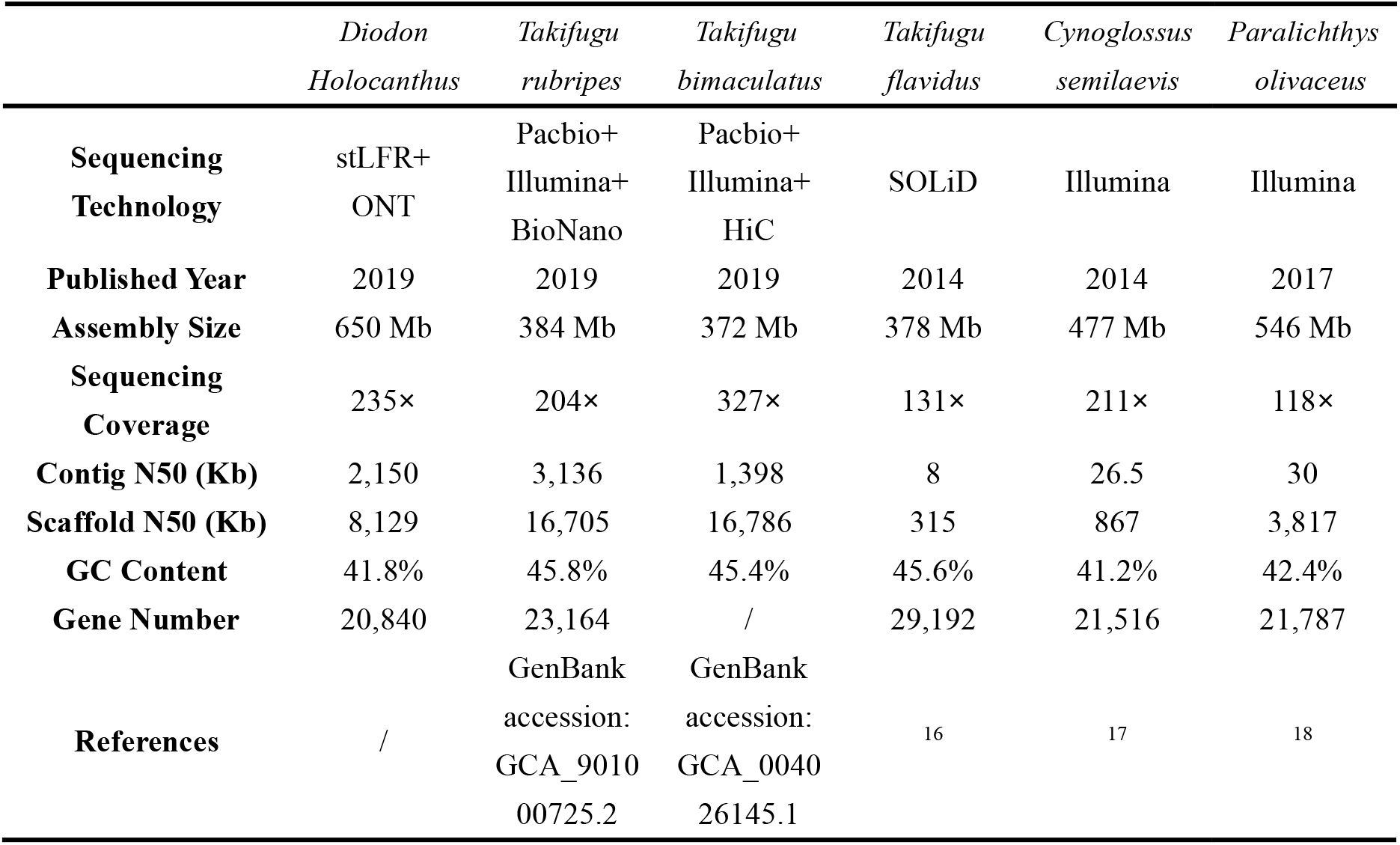
Summary statistics of recently published Tetraodontiforme genomes and relative species.

**Table 2.**
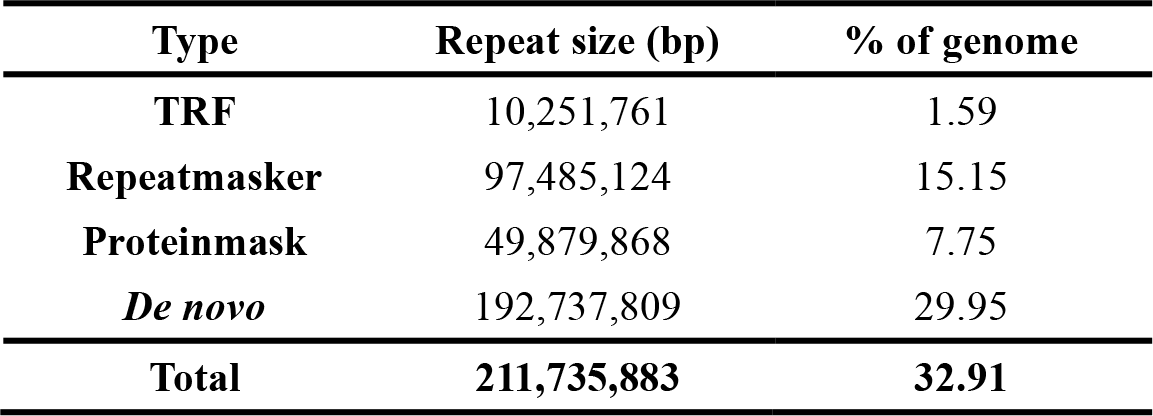
Prediction of repeat elements in the *Diodon Holocanthus* genome.

**Table 3.**
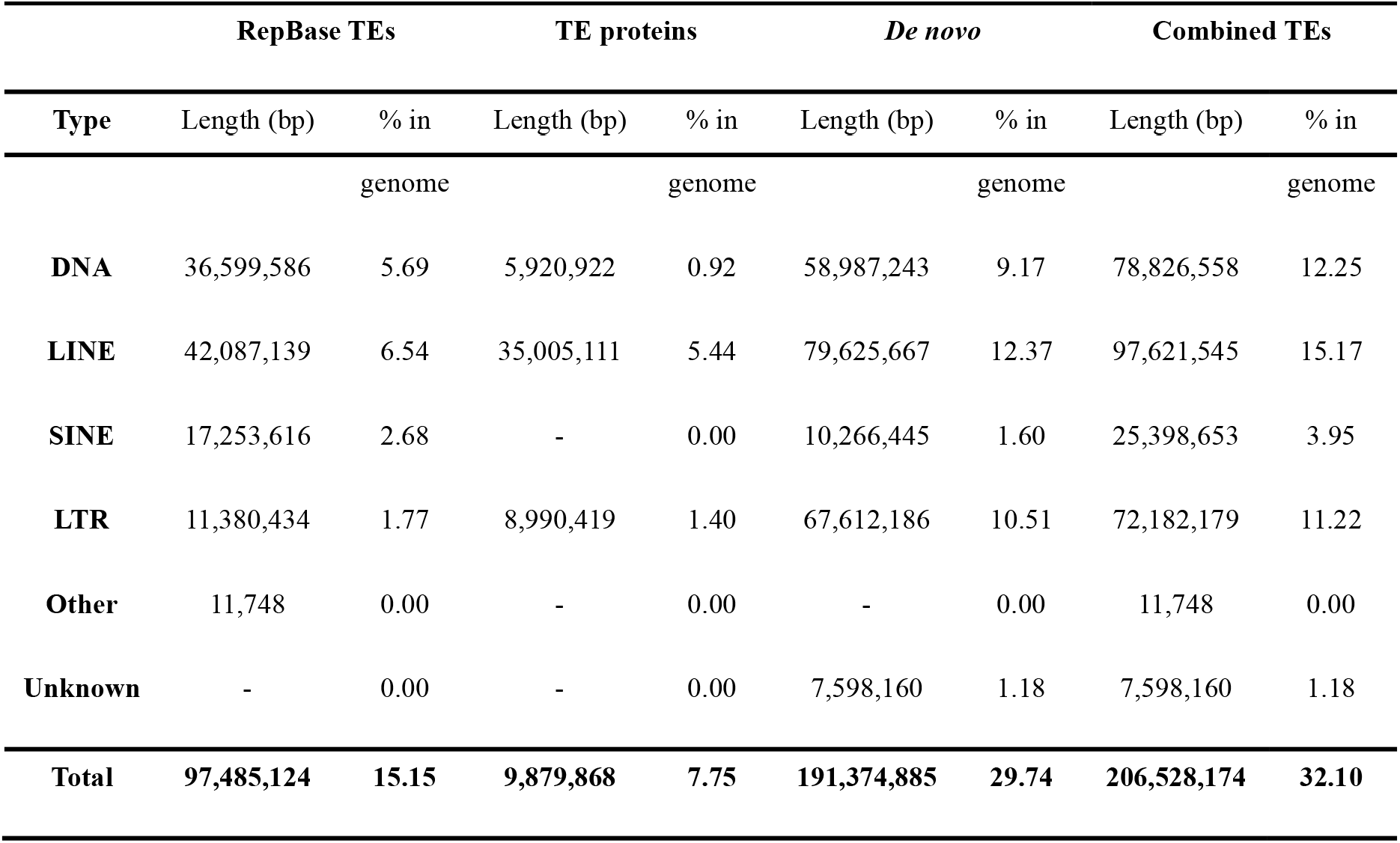
Statistics of repeat elements in the *Diodon Holocanthus* genome.

### Genome annotation

Structural annotation of genes. Two methods (homology-based and*ab initio* predictions) were used to predict spiny porcupinefish genes. In the homology-based method, the protein repertoires of *Astatotilapia calliptera*, *Maylandia zebra*, *Oreochromis niloticus* and *Pundamilia nyererei* were downloaded from the NCBI database and mapped onto the spiny porcupinefish genome using BLAT (version 0.36) ^24^ and GeneWise (version 2.4.1) ^25^ to define gene models. For the *ab initio* prediction approach, Augustus ^26^, GlimmerHMM ^27^, and GENSCAN ^28^ were applied to predict the coding regions of genes using *Danio rerio* as the model species of Hidden Markov Model method. In total, 20,840 non-redundant protein-coding genes were annotated in the spiny porcupinefish genome by combining the different evidences using EVidenceModeler (version 1.1.1) ^29^ (Table 4). The average length was 13,736.38 bp, with an average of 9.71 exons. The average length of coding sequences, exons and introns were 1,677.23bp, 172.80 bp and 1385.12 bp, respectively, similar to that of the other released fish genomes, such as *Astatotilapia calliptera*, *Maylandia zebra*, *Oreochromis niloticus* and *Pundamilia*.

**Table 4.** Comparison on gene structures of annotated gene models of the *Diodon Holocanthus* with other species.

| Gene set | # of genes | CDS+intron length (avg) | CDS length (avg) | Exon length (avg) | Intron length (avg) | Exons per gene (avg) |
| --- | --- | --- | --- | --- | --- | --- |
| *Diodon Holocanthus* | 20,840 | 13736.38 | 1677.23 | 172.8 | 1385.12 | 9.71 |
| *Danio_rerio* | 25,619 | 25207.59 | 1642.64 | 174.39 | 2798.97 | 9.42 |
| *Oryzias_latipes* | 19,699 | 12145.58 | 1515.82 | 147.82 | 1148.61 | 10.25 |
| *Takifugu_rubripes* | 18,523 | 7492.75 | 1693.53 | 152.61 | 574.33 | 11.1 |

Several databases including TrEMBL ^30^, SWISS-PROT ^31^ and InterPro ^32^ were used to search homologs to detect functions of the annotated protein-coding genes. 18,214 (87.40%), 17,015 (81.65%), 4,812 (23.09%) of protein-coding genes were found to present their homologous alignments in the three databases, respectively. We note that the remaining 2,559 (12.28%) protein-coding genes which cannot be identified and functionally annotated by existing database might be related to the specific characters of the *Diodon Holocanthus* genome, deserving further investigation.

## Data Records

Raw reads from BGISEQ-500 sequencing are deposited in the CNGB Nucleotide Sequence Archive (CNSA) with accession number CNP0000691 (https://db.cngb.org/cnsa). Data Citation 1: CNGB Nucleotide Sequence Archive CNP0000691).

## Author contributions

XXX

## Acknowledgements

This work was supported by the special funding of “Blue granary” scientific and technological innovation of China (2018YFD0900301-05). We also appreciate the support and help from the Fish10K project. The work also received the technical support from China National Gene Bank.

