## Supplementary_Information for "Draft genome of a porcupinefish, *Diodon Holocanthus*"

| Platform | Metric | Value |
|---|---|---|
| stLFR co-barcoded reads | Insert Size | 250 |
| | Read length (bp) | 100 |
| | Total data (Gb) | 154.3 |
| | Sequence coverage (×) | 219.8 |
| MinION long reads | # of Reads | 655,108 |
| | Average length (bp) | 17,378 |
| | Total data (Gb) | 11.4 |
| | Sequence coverage (×) | 15.9 |

**Table S1. Descriptive metrics of the input sequence data for the *de novo* genome assembly.**

| Kmer | Kmer Depth | Kmer Number | Estimated genome size (Mb) | Heterozygous Rate (%) | Repeat Rate (%) |
|---|---|---|---|---|---|
| 17 | 110 | 78,590,182,272 | 701.96 | 0.76 | 36.35 |

**Table S2. Estimation of the genome size, heterozygous and repeat rate using K-mer analysis.**

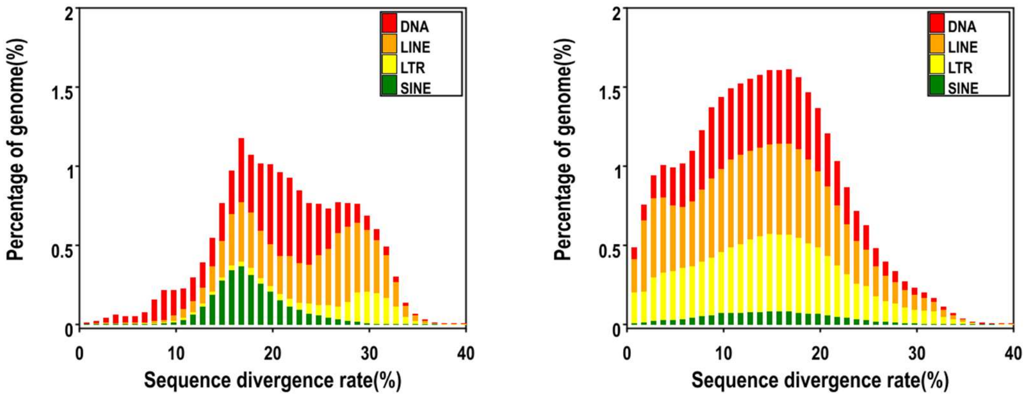


**Figure S1. Statistics of repeat elements in the *Diodon Holocanthus* genome.**

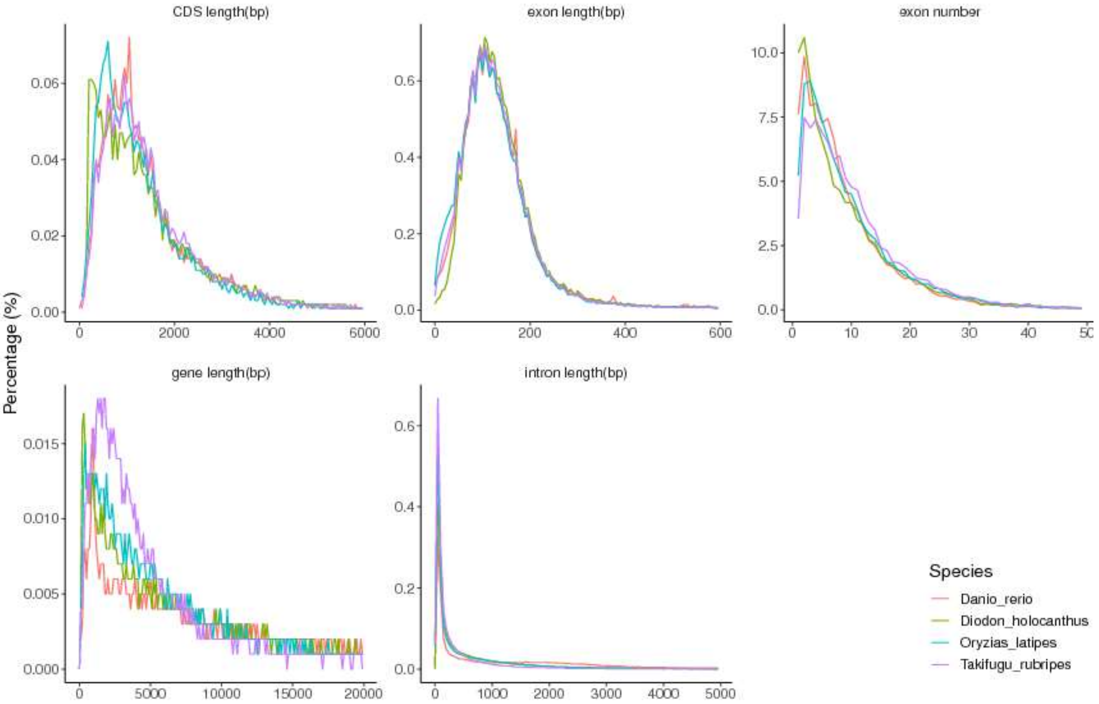


**Figure S2. Length distribution comparison on total gene, CDS, exon, and intron of annotated gene models of the *Diodon Holocanthus* with other species. Length distribution of total genes (A), CDS (B), exon (C), and intron (D) were compared to those of *Danio rerio, Oryzias latipes and Takifugu rubripes*.**

| Values | Total | Swissprot-Annotated | TrEMBL-Annotated | Interpro-Annotated | Overall |
|---|---|---|---|---|---|
| Number | 20,840 | 17,015 | 18,214 | 4,812 | 18,281 |
| Percentage | 100% | 81.65% | 87.40% | 23.09% | 87.72% |

**Table S3. The number of genes with homology or functional classification for *Diodon Holocanthus.***
